## Supplemental information for "Synaptotagmin-1-dependent phasic axonal dopamine release is dispensable for basic motor behaviors in mice"

**Supplemental Table 1:** *primers used for RT-qPCR quantifications.*

| Gene | UPL Probe | Oligo FWD | Oligo REV | RefSeq of isoform detected | Efficiency |
|---|---|---|---|---|---|
| ***GAPDH*** | 80 | agccacatcgctcagacac | gcccaatacgaccaaatcc | NM_008084.2 | 91% |
| ***th*** | 15 | cccaagggcttcagaagag | gggcatcctcgatgagact | NM_009377 | 106% |
| ***Slc18a2*** | 25 | caactttggagttggttttgc | ccaccaggtagcccatgata | NM_172523.3 | 93% |
| ***Slc6a3*** | 16 | ggaggcttacaggacctcaa | gagcccgaggaagtctgttt | NM_010020.3 | 95% |
| ***Syt1*** | 63 | gggaagaccatgaaggatca | tgaagctttcccagtttctcc | NM_009306.3, NM_001252341.1, NM_001252342.1 | 90% |
| ***Syt4*** | 25 | cccagaaaacctaagtagcaaaaa | agagcggctttacccttcac | NM_009308.3 | 90% |
| ***Syt7*** | 6 | gacgccacacgatgagtct | ctcagaacccggggagag | NM_018801.3, NM_173067.3, NM_173068.2 | 103% |
| ***Syt11*** | 27 | ggggtgtgtgctgtaaaagg | gtgctcacggggatatctcta | NM_018804.3 | 90% |
| ***Doc2b*** | 1 | ggctgatccctacgtcaaac | ccgaagagtttttgttctgagc | NM_007873.3 | 98% |

**Supplemental figure and legends**

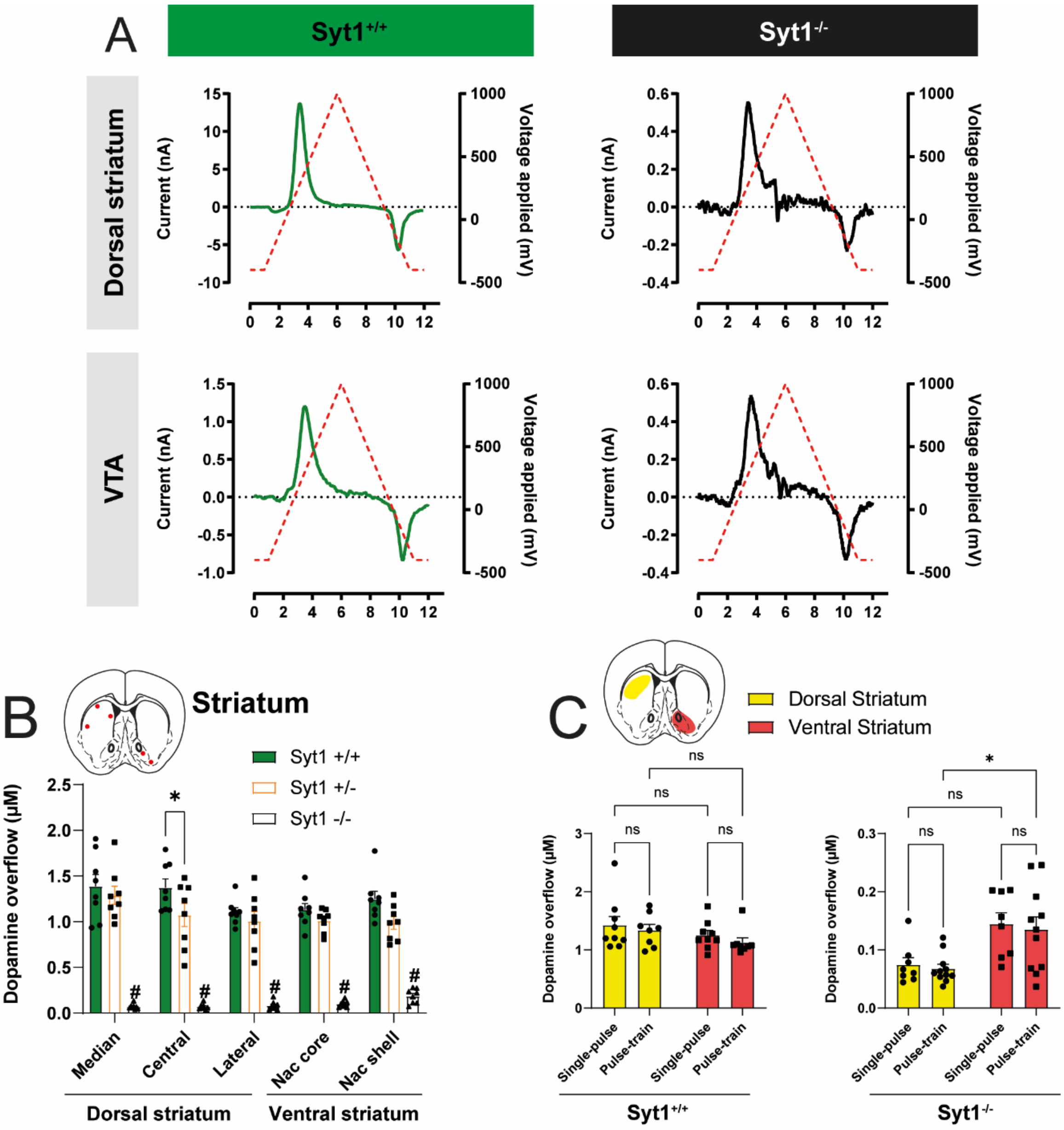


***Figure S1: FSCV recordings in Syt1 cKO$^{DA}$ mice. (A)** Representative voltammograms obtained in the dorsal striatum and VTA from Syt1$^{+/+}$ and Syt1$^{-/-}$ mice using a voltage ramp of -400 to 1000 mV vs Ag/AgCl at 300 V/s with a 100 ms sampling interval. **(B)** Regional sampling (single pulse, 1 ms, 400 µA) of DA release in Syt1$^{+/+}$ (n = 18 slices/9 mice), Syt$^{+/-}$ (n = 16/8) and Syt1$^{-/-}$ mice (n = 16/8) in the dorsal striatum (median, central and lateral)*

*and ventral striatum (Nac core and shell). Statistical analysis was carried out by a 2-way ANOVA followed by a Tukey test.* ***(C)*** *Quantification of peak amplitude measured in the dorsal (yellow) and ventral striatum (red) of* $Syt^{+/+}$ *(left) and* $Syt1^{-/-}$ *(right) mice using single pulse (n = 18/9 in* $Syt^{+/+}$ *and n = 16/8 in* $Syt1^{-/-}$*) or pulse-train stimulation (n = 16/8 in* $Syt^{+/+}$ *and n = 22/11 in* $Syt1^{-/-}$*). Statistical analysis was carried out by a 2-way ANOVA followed by a Šidák's correction. Error bars represent* ± *S.E.M. (ns, non-significant; *,* $p < 0.05$*; **,* $p < 0.01$*; ***,* $p < 0.001$*; #,* $p < 0.0001$*).*

Fig. S2

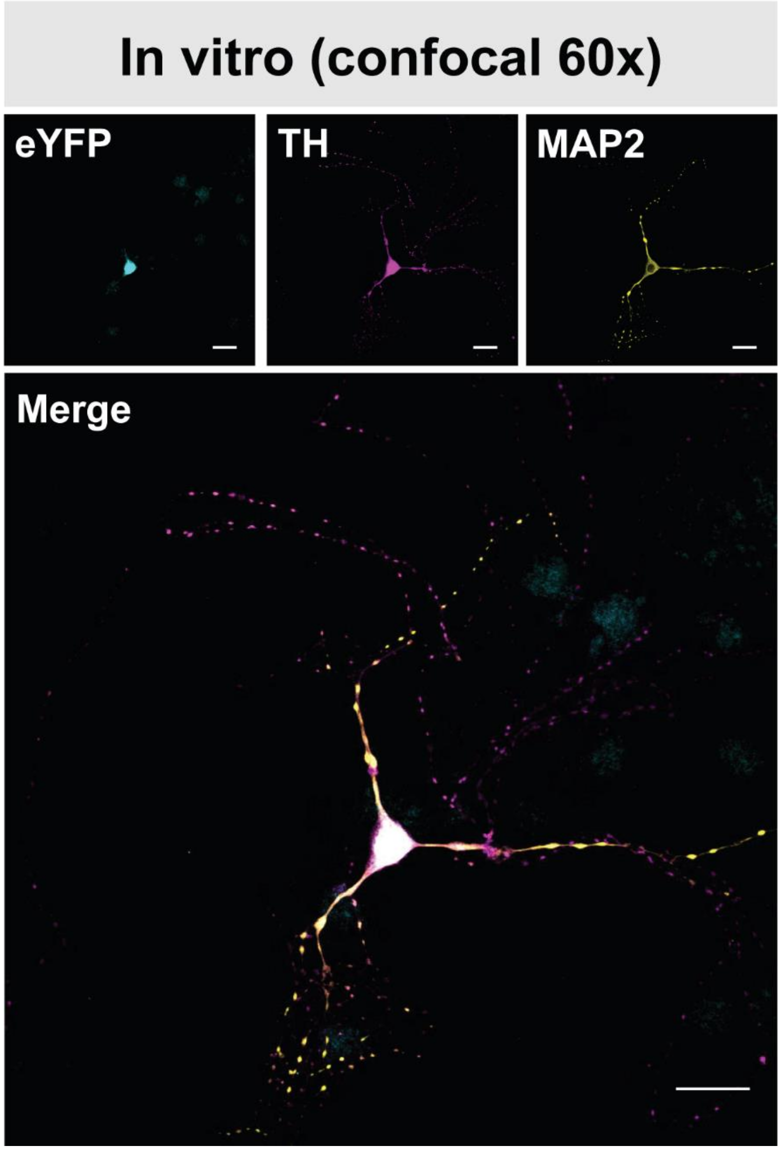


***Figure S2: Somatodendritic subcellular localization of ChR2-Kv in cultured dopamine neurons.*** *Immunocytochemistry of one representative TH positive DA neuron infected with AAV2/5-hsyn-DIO-ChR2-eYFP-Kv shows the expression of ChR2-Kv (eYFP) within the soma and proximal dendrite (TH$^{+}$/MAP2$^{+}$), but a lack of expression in the axonal compartment (TH$^{+}$/MAP2$^{-}$ processes), with confocal microscopy at 60x (scale bar = 20 µm). A total of 2 different coverslips of cultured primary neurons were analyzed by IHC.*

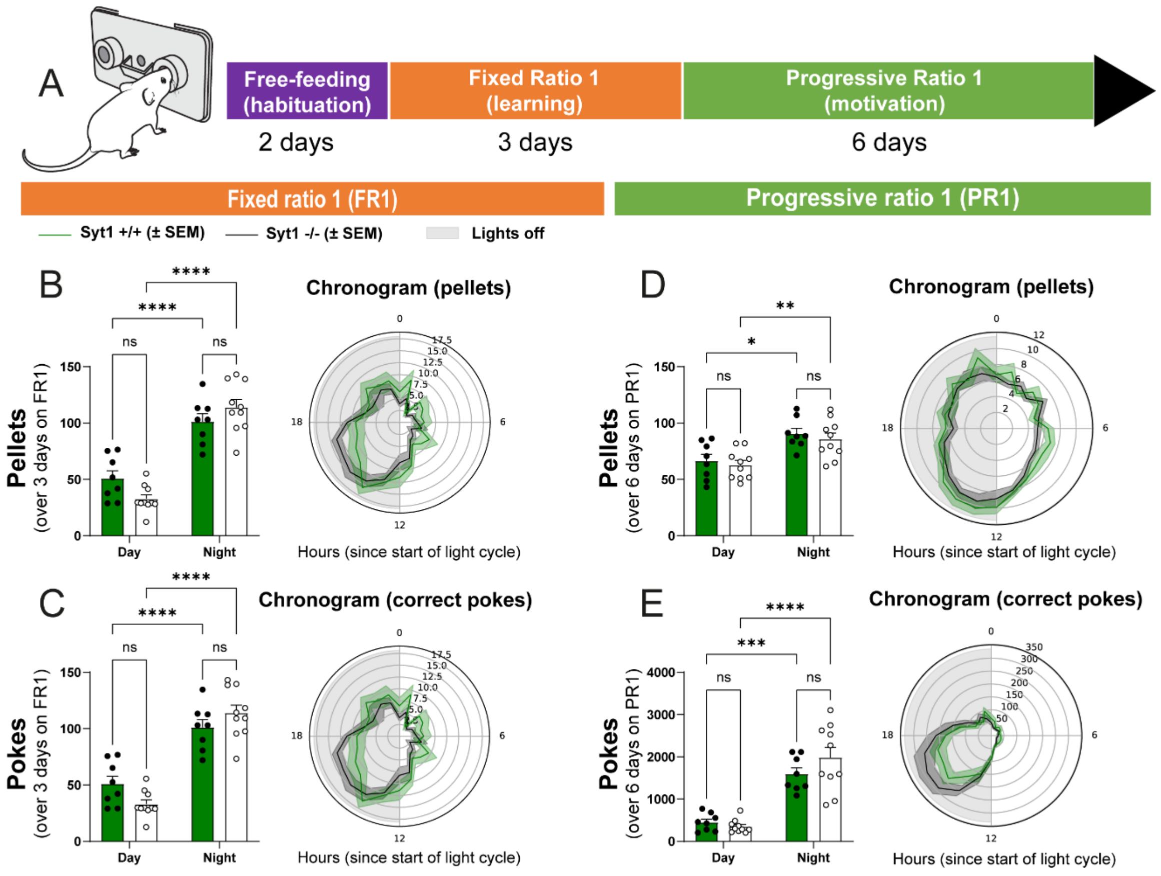


***Figure S3: Syt1 is dispensable for motivation to work for food. (A)** Schematic representation of the operant food-rewarded nose-poking protocol used. The diagram is modified from https://github.com/KravitzLabDevices/FED3/blob/main/photos/mouse_feeder.svg. **(B-C)** number of pellets earned **(B)** and pokes made **(C)** by Syt1$^{+/+}$ (green, n = 8) and Syt1$^{-/-}$ (black, n = 10) mice during night or day cycle measured with "FED3" feeding devices, over 3 days in a fixed ratio (FR) 1 paradigm (1 pellet for 1 poke). **(D-E)** number of pellets earned **(D)** and pokes made **(E)** by Syt1$^{+/+}$ (n = 8) and Syt1$^{-/-}$ (n = 10) mice during night or day cycle measured over 6 days in a progressive ratio (PR) 1 paradigm (the nose-poking requirement began on FR1, increased by 1 poke each time a pellet was earned and was reset to FR1 if no poking was done on either the active or inactive port for 30 min). For*

*each experiment, mice were placed in a 12h/12 light/dark cycle. Chronograms of pellets eaten and pokes on the active port during each period is presented on the right side of each graph. Statistical analyses were carried out by 2-way ANOVAs followed by Tuckey test. Error bars represent ± S.E.M (ns, non-significant; *, $p < 0.05$; **, $p < 0.01$; ***, $p < 0.001$; ****, $p < 0.0001$).*

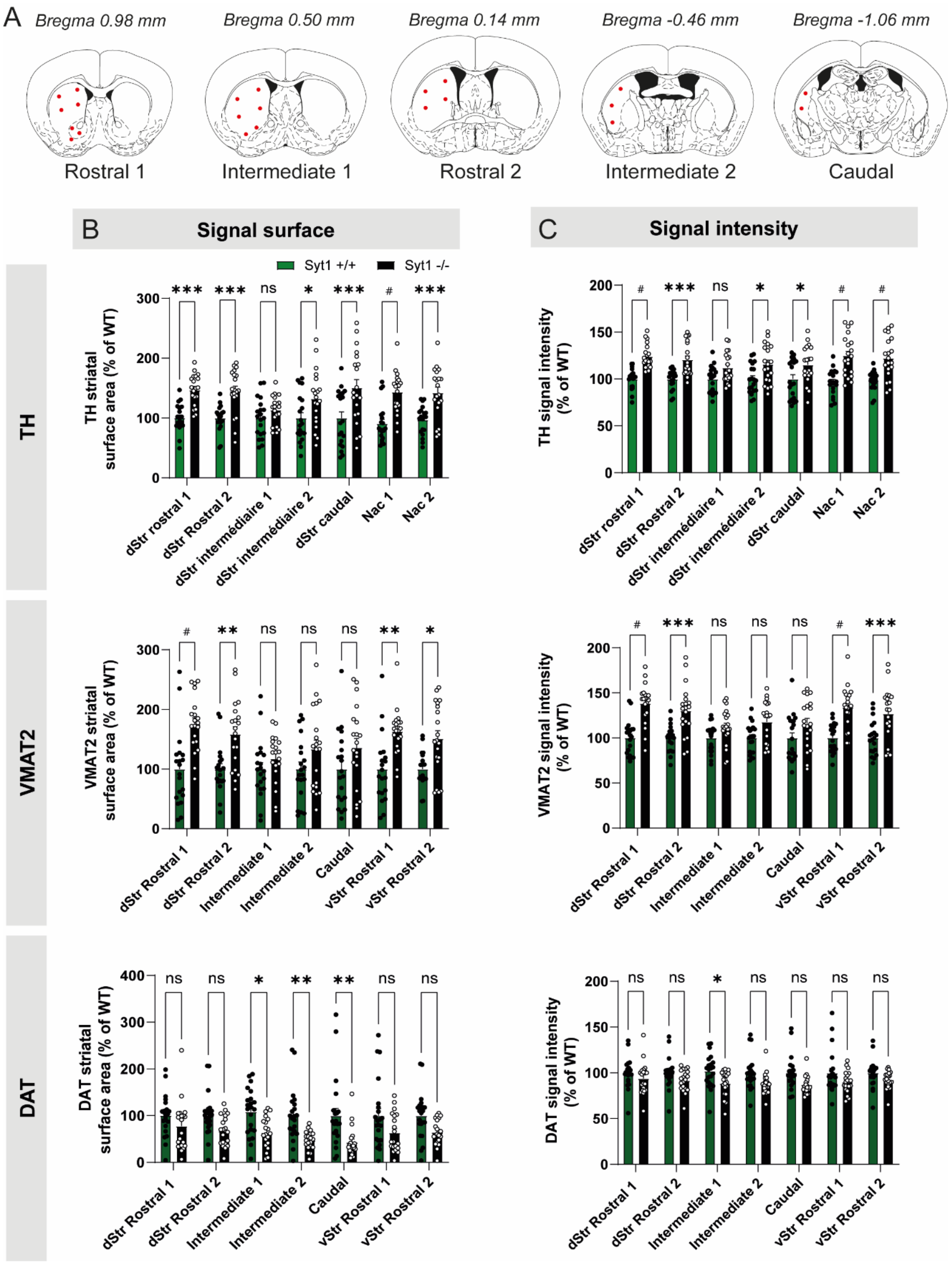


***Figure S4: Regional adaptations of the dopaminergic system of Syt1 cKO$^{DA}$ mice. (A)***

*Schematic representation of striatal slices used for immunohistochemistry characterization. Surface and intensity for each signal were measured in a series of 5 different striatal slices ranging from bregma +0.98 to bregma -1.06 mm, with a total of 22 different areas for each hemisphere.* ***(B)*** *Quantification of signal surface (% of WT) for TH, VMAT2, DAT and 5-HT in the different striatal regions examined: "rostral 1 and 2, intermediate 1 and 2 and caudal (n = 20 hemispheres/10 mice for both genotypes). "Nac 1 and 2" referring to the ventral striatum (nucleus accumbens) from the rostral 1 and 2 slices respectively.* ***(C)*** *Same for signal intensity of each immunostaining (n = 20 hemispheres/10 mice for both genotypes). Statistical analysis was carried out by 2-way ANOVAs followed by Šidák's corrections. Error bars represent ± S.E.M. (ns, non-significant; *, $p < 0.05$; **, $p < 0.01$; ***, $p < 0.001$; #, $p < 0.0001$).*

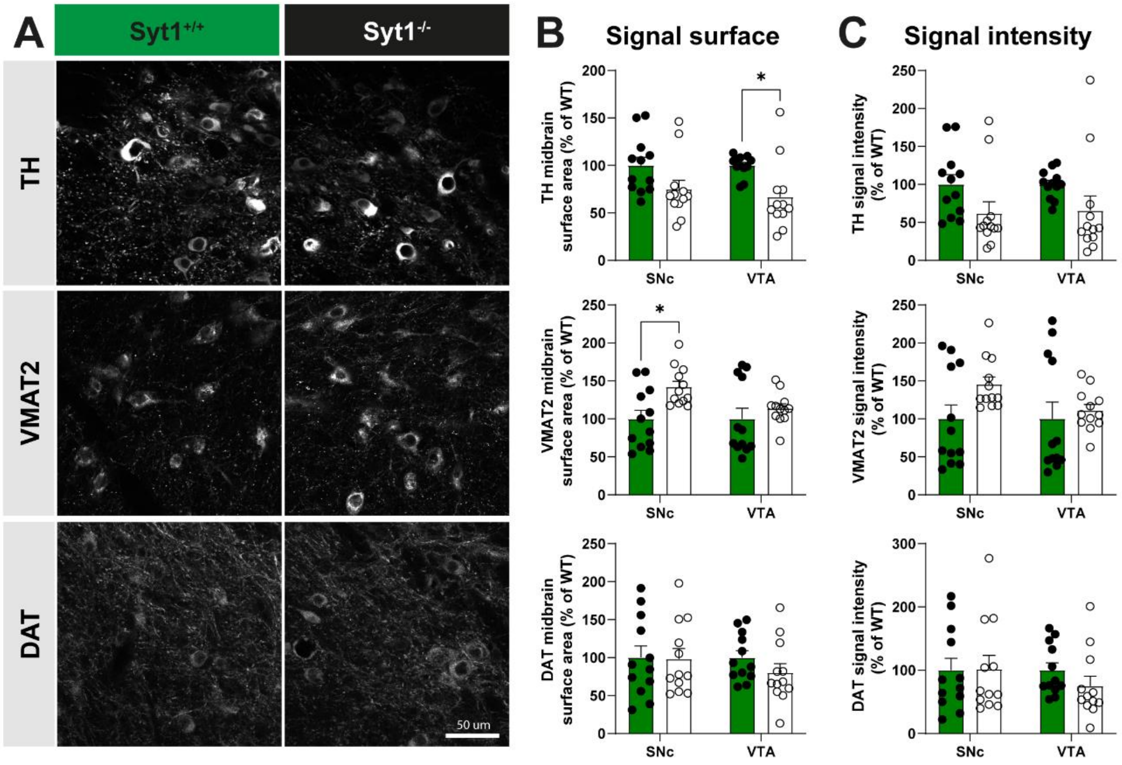


***Figure S5: Adaptations of DA innervation in the mesencephalon of Syt1 cKO$^{DA}$ mice.***

***(A)*** *Immunohistochemistry of mesencephalic slices from 10-12-week-old Syt1$^{+/+}$ and Syt1$^{-/-}$ mice (60x confocal) using (from top to bottom): tyrosine hydroxylase (TH), vesicular monoamine transporter 2 (VMAT2) and dopamine transporter (DAT) immunostainings. Scale bar = 50µm.* ***(B)*** *Quantification of each signal surface (% of WT) in the SNc and VTA of Syt1$^{+/+}$ and Syt1$^{-/-}$ mice (n = 12 hemispheres/6 mice).* ***(C)*** *Same with signal intensity (n = 12 hemispheres/6 mice for both genotypes). Statistical analysis was carried out by 2-way ANOVAs followed by Šidák's corrections. Error bars represent ± S.E.M. (ns, non-significant; *, $p < 0.05$; **, $p < 0.01$; ***, $p < 0.001$; ****, $p < 0.0001$).*

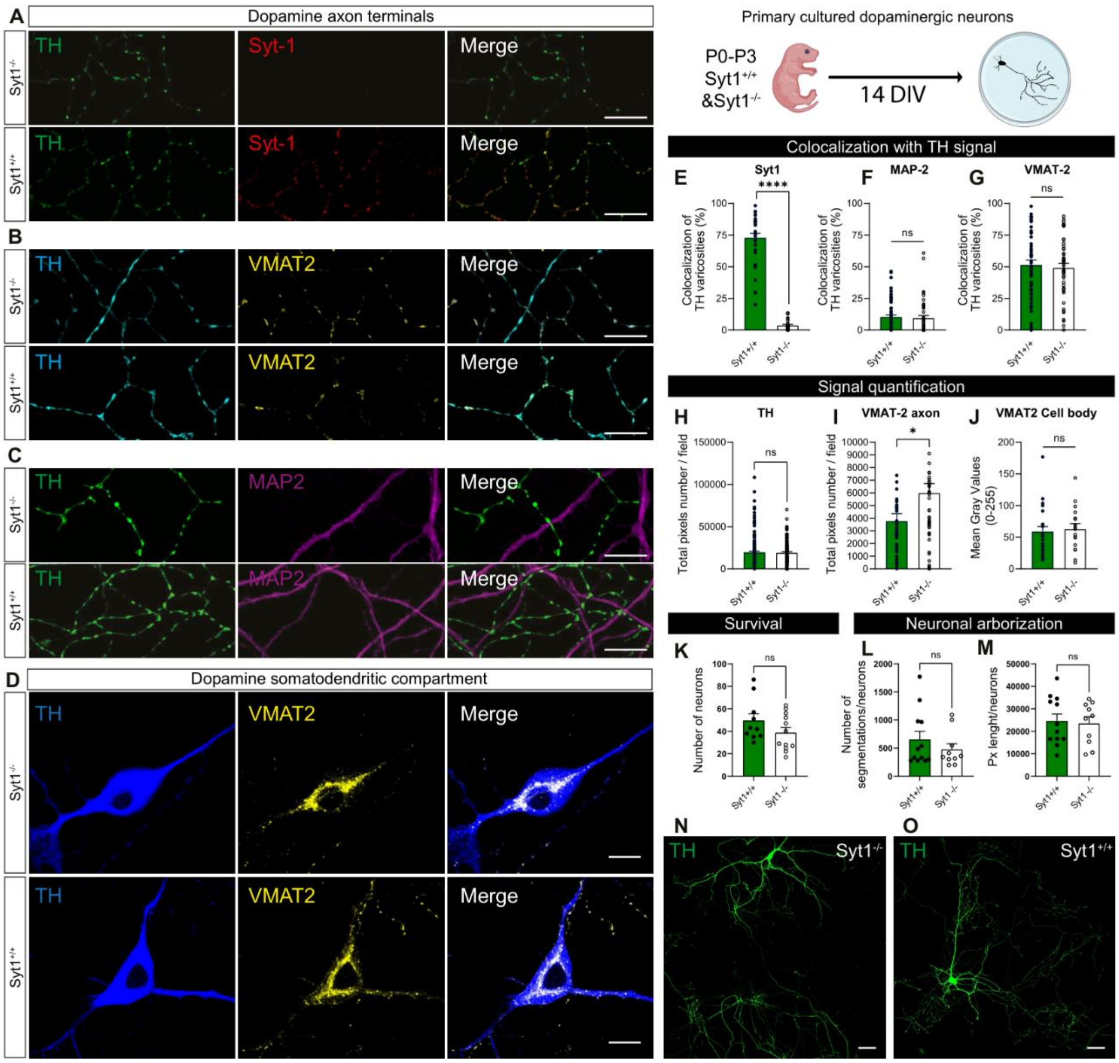


***Figure S6: No significant changes in the overall morphological development of cultured DA neurons.*** *Primary DA neurons from VTA or SNc were cultured from Syt1$^{-/-}$ or Syt1$^{+/+}$ P0-P3 pups and used for immunocytochemistry (ICC) at 14 days in vitro (DIV).* ***(A)*** *Immunocytochemistry of cultures prepared from Syt1$^{+/+}$ and Syt1$^{-/-}$ mice showing colocalization of Syt1 signal with TH$^{+}$ axonal varicosities (confocal 60x, scale bar = 10 µm).* ***(B)*** *Same with VMAT2 antibody (confocal 60x, scale bar = 10 µm).* ***(C)*** *Synaptic contacts of TH$^{+}$ varicosities on MAP2$^{+}$/TH$^{-}$ dendrites (confocal 60x, scale bar = 10 µm).* ***(D)*** *Colocalization of VMAT2 signal with TH$^{+}$ soma and dendrites (scale bar = 5 µm).* ***(E-***

***G)*** *Quantifications of Syt1 (n = 29 fields for Syt1*$^{-/-}$ *and 35 for Syt1*$^{+/+}$*)* ***(E)****, MAP2 (n = 47 fields for Syt1*$^{-/-}$ *and 64 for Syt1*$^{+/+}$*)* ***(F)*** *and VMAT2 (n = 50 fields for Syt1*$^{-/-}$ *and 55 for Syt1*$^{+/+}$*)* ***(G)*** *percentage of colocalization with TH*$^{+}$ *varicosities.* ***(H-J)*** *Quantification of TH (n = 138 fields for Syt1*$^{-/-}$ *and 166 for Syt1*$^{+/+}$*)* ***(H)*** *and axonal (n = 43 fields for Syt1*$^{-/-}$ *and 45 for Syt1*$^{+/+}$*)* ***(I)*** *and somatodendritic (n = 17 fields for Syt1*$^{-/-}$ *and 22 for Syt1*$^{+/+}$*)* ***(J)*** *VMAT2 signals.* ***(K)*** *Evaluation of cultured Syt1*$^{+/+}$ *and Syt1*$^{-/-}$ *DA neuron survival, assessed by their number/coverslips at 14 DIV (n = 10 coverslips Syt1*$^{+/+}$ *and 12 Syt1*$^{-/-}$*).* ***(L-M)*** *Evaluation of cultured Syt1*$^{+/+}$ *and Syt1*$^{-/-}$ *DA neuron development, assessed by the number of TH*$^{+}$ *processes* ***(L)*** *and their length* ***(M)*** *(n = 10 Syt1*$^{+/+}$ *and 12 Syt1*$^{-/-}$ *neurons).* ***(N-O)*** *Representative TH*$^{+}$ *DA neurons at 14 DIV from Syt1*$^{-/-}$ *(N) and Syt1*$^{+/+}$ *pups (confocal 20x, scale bar = 50 µm). Statistical analysis was carried out by two-tailed unpaired t-tests. Error bars represent ± S.E.M. (ns, non-significant; *, $p < 0.05$; **, $p < 0.01$; ***, $p < 0.001$; ****, $p < 0.0001$).*

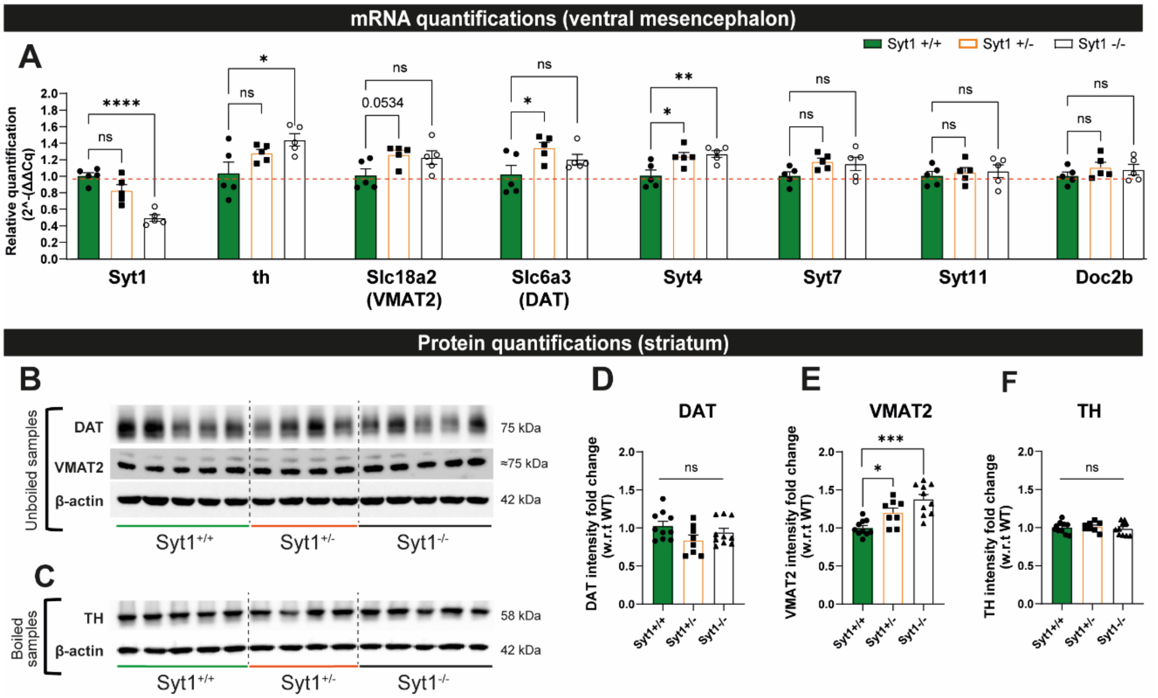


***Figure S7: Gene expression and protein levels in Syt1 cKO$^{DA}$ mice.*** ***(A)*** *Relative changes of mRNA levels measured by RT-qPCR, from microdissected ventral mesencephalon containing the SN/VTA of adult Syt1$^{+/+}$, Syt1$^{+/-}$ and Syt1$^{-/-}$ mice (n = 5 per genotype). Ct values (mean of duplicate repeats) of th, Slc18a2 (VMAT2), Slc6a3 (DAT), Syt1, 4, 7 and 11 mRNA levels were normalized to the Ct value of GAPDH in the same samples. Statistical analysis was carried out by 1-way ANOVA with Dunnett's tests.* ***(B)*** *Representative western blots illustrating either DAT, VMAT2 and β-actin from total striatum unboiled homogenates of adult Syt1$^{+/+}$, Syt1$^{+/-}$ and Syt1$^{-/-}$ mice.* ***(C)*** *Same for TH and β-actin in boiled samples.* ***(D-F)*** *Immunoblot quantifying relative protein levels (fold change compared to controls) for DAT* ***(D)****, VMAT2* ***(E)*** *and TH* ***(F)*** *(n = 10 Syt1$^{+/+}$ and Syt1$^{-/-}$ mice, n = 8 Syt1$^{+/-}$). The samples derive from the same set of experiments and the blots were processed in parallel. The results from different blots were pooled after normalizing to WT control (Syt1$^{+/+}$) samples. Statistical analysis was carried out by 1-way ANOVA with*

*Dunnett's tests (S7D and S7F) and Brown-Forsythe ANOVA with Dunnett's T3 test (S7E). Error bars represent ± S.E.M. (ns, non-significant; *, $p < 0.05$; **, $p < 0.01$; ***, $p < 0.001$; ****, $p < 0.0001$).*

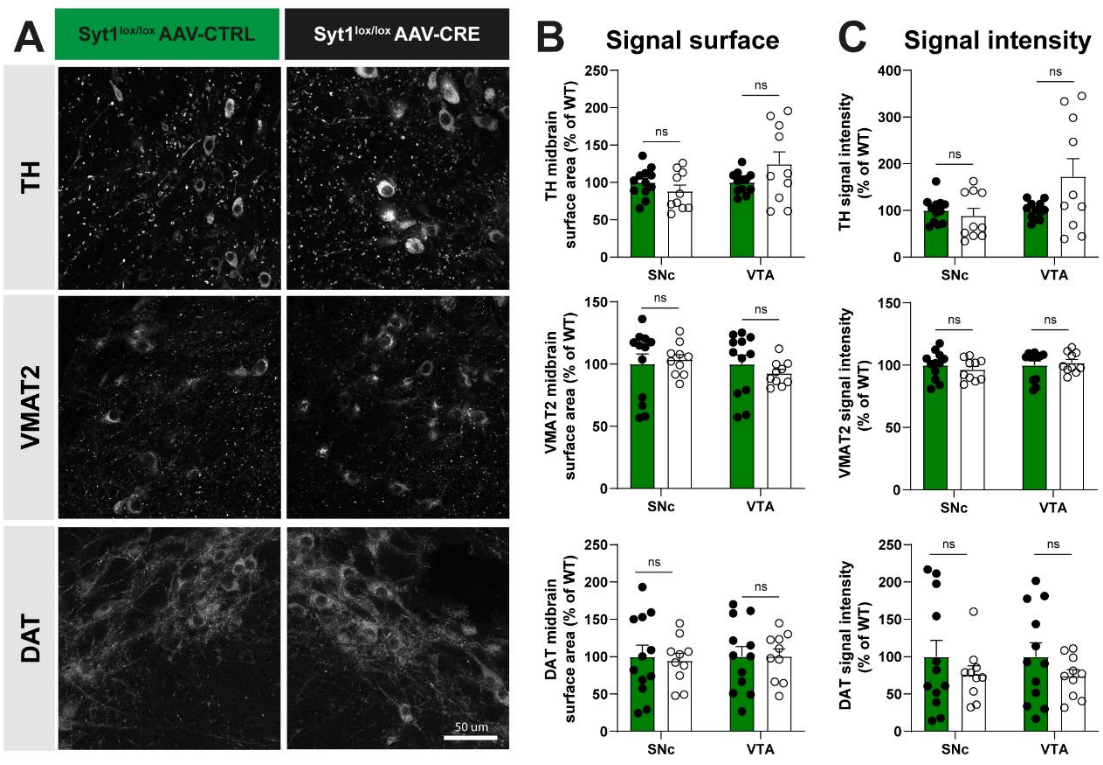


***Figure S8: Adaptations of DA innervation in the mesencephalon of mice with acute Syt1 deletion. (A)*** *Immunohistochemistry of mesencephalic slices from 9–10-week-old floxed-Syt1 mice injected at 6-7 weeks with an AAV9-TH-cre-myc-2A-fusion-red and control virus (60x confocal) using (from top to bottom): tyrosine hydroxylase (TH), vesicular monoamine transporter 2 (VMAT2) and dopamine transporter (DAT) immunostainings. Scale bar = 50 µm.* ***(B)*** *Quantification of each signal surface (% of WT) in the SNc and VTA of Syt1*$^{+/+}$ *and Syt1*$^{-/-}$ *mice (n = 12 hemispheres/6 Syt1*$^{+/+}$ *mice and n = 10/5 Syt1*$^{-/-}$ *mice).* ***(C)*** *Same with signal intensity (n = 12 hemispheres/6 Syt1*$^{+/+}$ *mice and n = 10/5 Syt1*$^{-/-}$ *mice). Statistical analysis was carried out by 2-way ANOVAs followed by Šidák's corrections. Error bars represent ± S.E.M. (ns, non-significant; *, $p < 0.05$; **, $p < 0.01$; ***, $p < 0.001$; ****, $p < 0.0001$).*
